# A new Paleogene fossil and a new dataset for waterfowl (Aves: Anseriformes) clarify phylogeny, ecological evolution, and avian evolution at the K-Pg Boundary

**DOI:** 10.1101/2022.11.23.517648

**Authors:** Grace Musser, Julia A. Clarke

## Abstract

Despite making up one of the most ecologically diverse groups of living birds, comprising soaring, diving and giant flightless taxa, the evolutionary relationships and ecological evolution of Anseriformes (waterfowl) remain unresolved. Although Anseriformes have a comparatively rich, global Cretaceous and Paleogene fossil record, morphological datasets for this group that include extinct taxa report conflicting relationships for all known extinct taxa. Correct placement of extinct taxa is necessary to understand whether ancestral anseriform feeding ecology was more terrestrial or one of a set of diverse aquatic ecologies and to better understand avian evolution around the K-T boundary. Here, we present a new morphological dataset for Anseriformes that includes more extant and extinct taxa than any previous anseriform-focused dataset and describe a new anseriform species from the early Eocene Green River Formation of North America. The new taxon has a mediolaterally narrow bill which is not known in any previously described anseriform fossils other than portions of the pseudotoothed Pelagornithidae. The matrix created to assess the placement of this taxon comprises 41 taxa and 719 discrete morphological characters describing skeletal morphology, musculature, syringeal morphology, ecology, and behavior. We additionally combine the morphological dataset with published sequences using Bayesian methods and perform ancestral state reconstruction for ecological and behavioral characters. We recover the new Eocene taxon as a stem anseranatid across all analyses, and find that the new taxon represents a novel ecology within known Anseriformes and the Green River taxa. Results indicate that Anseriformes were likely ancestrally aquatic herbivores with rhamphothecal lamellae and provide insight into avian evolution during and following the K-Pg mass extinction.

## Introduction

Although extant Anseriformes may appear to occupy a relatively narrow range of dominantly aquatic ecologies, both extinct and extant taxa show a broad range of locomotor and feeding modes. Extinct ecotypes exhibited in the comparatively rich Cretaceous-Paleogene fossil record of this clade include the terrestrial, giant flightless birds *Diatryma* Cope 1876 and *Gastornis* Hébert 1855 (*vide* Prévost 1855) with dorsoventrally broad beaks and debated diets (Witmer and Rose, 1991; Andors, 1992); the marine albatross-like Pelagornithidae with pseudotooth projections for piscivorous diets (Mayr and Rubilar-Rogers, 2010; Ksepka, 2014; Mayr et al., 2021); and the more aquatic, duck-like species such as the wide-billed, but long-legged wader *Presbyornis* Wetmore 1926 (Livezey, 1997; Ericson, 1997). Extant Anseriformes include terrestrial and primarily herbivorous grazers of the Anhimidae; *Anseranas*, an aquatic surface swimmer and grazer that is primarily herbivorous; and a variety of ecotypes within Anatidae (Delacour and Mayr, 1945; Livezey, 1986). Within Anatidae, taxa possess both herbivorous and omnivorous diets with various degrees of specialization; different filter feeding modes comprising grazing, mixed feeders, and diving graspers; and a variety of swimming ecologies that comprise a terrestrial ecology, foot propelled swimming, wing propelled swimming, surface swimming, plunging, and foot and wing propelled swimming (Li and Clarke, 2015; Olsen, 2015; Olsen, 2017). Diving, as either an escape behavior or in feeding, occurs throughout Antatidae but not in Anhimidae and possibly not in *Anseranas* (Johnsgard, 1962; Todd, 1979; Livezey, 1986).

Despite the ecological breadth of anseriform ecologies and an abundance of recovered extinct taxa, important questions remain regarding the phylogeny and ecological and behavioral evolution of Anseriformes. It is debated when within stem or crown Anseriformes they evolve more aquatic ecologies, especially as vestigial rhamphothecal lamellae and pedal webbing were proposed to be present in extant Anhimidae (Olson and Feduccia, 1980). It is similarly uncertain how herbivory and beak shape evolved within this group. Simulated trait evolution has supported two main patterns of diversification of beak shape and related diet in crown Anseriformes: either a single evolutionary trajectory or several independent and parallel transitions to a narrower, more herbivorous “goose-like” beak are estimated (Olsen, 2017). Phylogenetic placement of extinct anseriform taxa is necessary to inform evolution of these traits in both stem and crown.

Morphological analyses containing extinct taxa have resulted in drastically differing topologies (Livezey, 1997; Ericson, 1997; Clarke et al., 2005; Bourdon, 2005; Mayr, 2011; Louchart et al., 2013; Worthy et al., 2017; Tambussi et al,. 2019; Field et al., 2020) and specimens that represent or are referrable to several extinct taxa have not been included or fully captured by scorings in previous analyses, resulting in a higher level of missing data in these analyses. In addition, fine-grained molecular analyses of extant taxa have produced conflicting results within Anatidae (Sraml et al., 1996; Donne-Goussé et al., 2002; Sun et al., 2017). Placement of extinct taxa is critical as ancestral state reconstructions are optimized differently depending on variations in extinct taxon placement. Better understanding the evolutionary relationships and ecological evolution of Anseriformes and their stem lineages is also critical for better understanding early avian evolution and biogeography. New Paleogene fossils and robust placement of these and previously described taxa within a phylogenetic context is necessary to better understand the answers to these questions.

The predominantly lacustrine Green River Formation of North America preserves an exceptional snapshot of early Eocene diversity in North America (Grande, 2013), but has only produced two aquatic taxa to date: *Presbyornis*, an anseriform, and *Limnofregata* Olson 1977, likely an extinct relative of living frigate birds (Olson and Matsuoka, 2005; Stidham, 2015). An ibis-like taxon of uncertain ecology has also been recovered (Smith et al., 2013). Here we describe a new aquatic avian taxon from the early Eocene Fossil Butte Member (FBM; 51.97 ± 0.16 Ma; Smith et al., 2010) of the Green River Formation. The taxon was originally illustrated with a prospective referral by Storrs Olson to Heliornithidae (finfoots; Grande 2013); however, we recover the taxon as a member of Anseriformes. Known Paleogene Anseriformes have a wide, duck-like beak (i.e. *Presbyornis*, *Anatalavis* Olson and Parris 1987), narrow pseudothoothed or dorsoventrally broad beaks (eg. *Pelagornis* Lartet 1857 and *Diatryma*, respectively), or largely unknown beak morphology (i.e. *Conflicto antarcticus* Tambussi et al. 2019; *Vegavis iaai* Clarke et al. 2005; Clarke et al., 2016). In contrast, the new fossil presents a narrow bill that is most similar to the Anhimidae but differs from all extant Anseriformes. We identify this fossil as the holotype specimen of a new species. New x-ray computed tomography (CT) images allow recovery of previously hidden morphologies, revealing it to represent a new taxon and ecology for the Green River Formation. The specimen was found at a near-shore locality of the Fossil Butte Member of the Formation, locality H (Grande and Buchheim, 1994; Grande, 2013), where several lithornithid and neoavian fossils have been previously described (Grande, 2013; Nesbitt and Clarke, 2016).

We additionally present a new morphological dataset for extinct and extant Galloanseres which includes more examplars of extinct and extant Anseriformes than any previous matrix.

Creation of this dataset included reassessment of and scorings from many specimens of *Presbyornis* and *Telmabates antiquus* Howard 1955 that have and have not been included in previous studies. This included re-evaluation of all *Presbyornis* material housed in the Vertebrate Paleontology Collection of the Smithsonian National Museum of Natural History and all *Telmabates* material housed in the Vertebrate Paleontology Collection of the American Museum of Natural History. We additionally combine the new morphological matrix with molecular data for Bayesian analysis and perform ancestral state reconstruction of ecological and behavioral traits as well as principle component and linear discriminant analyses. Results allow new insights into outstanding issues of phylogeny, evolution and avian diversification.

### Institutional Abbreviations

AMNH, American Museum of Natural History, New York, NY, U.S.A.; FMNH, Field Museum of Natural History, Chicago, IL, U.S.A.; TMM, the Texas Memorial Museum, Austin, Texas, U.S.A.; USNM, National Museum of Natural History, Smithsonian Institution, Washington, D.C., U.S.A. Specimen numbers are presented in Table 1.

**Table 1.**
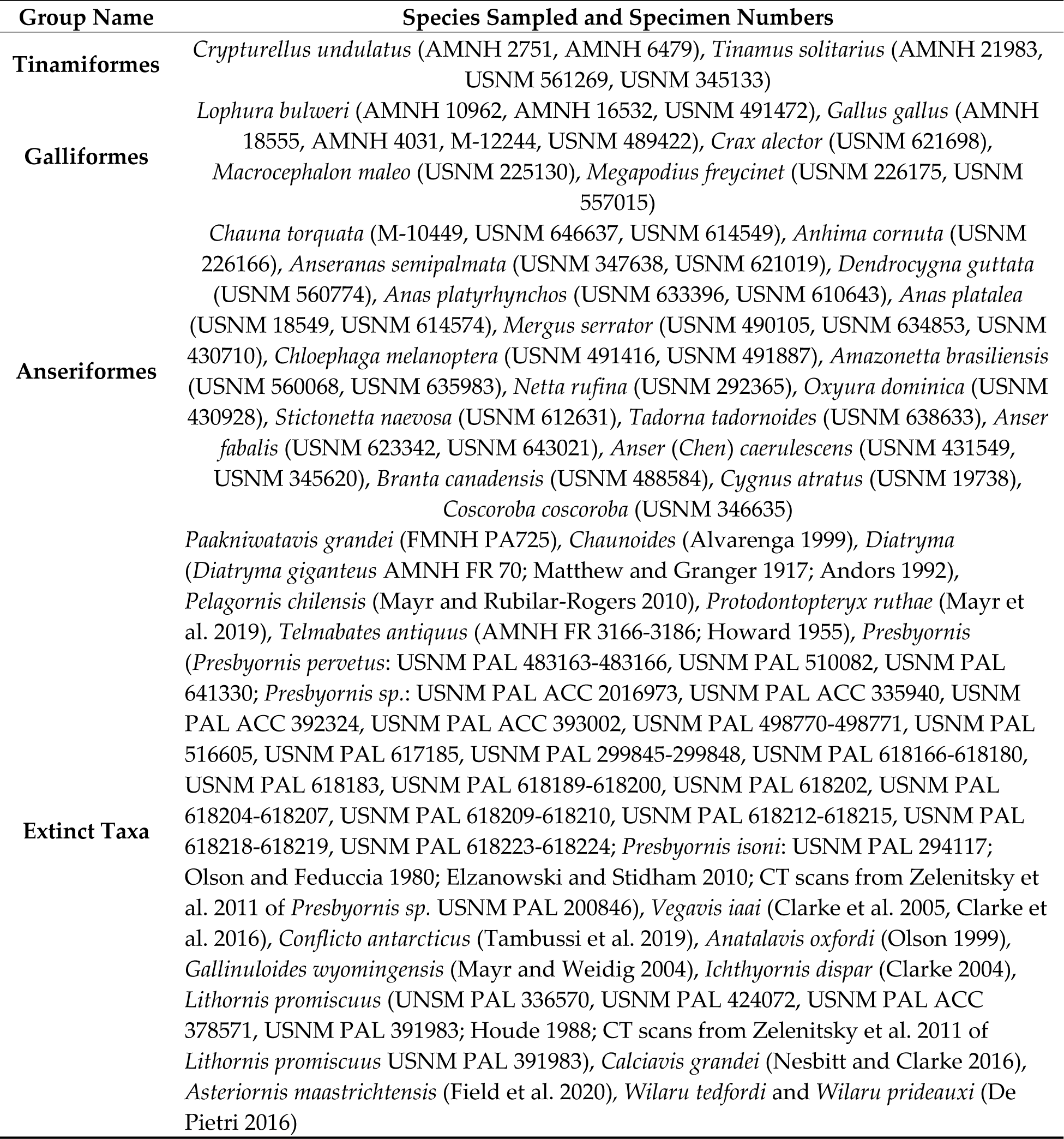
Specimen numbers of skeletal specimens used for comparison during fossil description and phylogenetic analyses.

| Group Name | Species Sampled and Specimen Numbers |
| --- | --- |
| **Tinamiformes** | *Crypturellus undulatus* (AMNH 2751, AMNH 6479), *Tinamus solitarius* (AMNH 21983, USNM 561269, USNM 345133) |
| **Galliformes** | *Lophura bulweri* (AMNH 10962, AMNH 16532, USNM 491472), *Gallus gallus* (AMNH 18555, AMNH 4031, M-12244, USNM 489422), *Crax alector* (USNM 621698), *Macrocephalon maleo* (USNM 225130), *Megapodius freycinet* (USNM 226175, USNM 557015) |
| **Anseriformes** | *Chauna torquata* (M-10449, USNM 646637, USNM 614549), *Anhima cornuta* (USNM 226166), *Anseranas semipalmata* (USNM 347638, USNM 621019), *Dendrocygna guttata* (USNM 560774), *Anas platyrhynchos* (USNM 633396, USNM 610643), *Anas platalea* (USNM 18549, USNM 614574), *Mergus serrator* (USNM 490105, USNM 634853, USNM 430710), *Chloephaga melanoptera* (USNM 491416, USNM 491887), *Amazonetta brasiliensis* (USNM 560068, USNM 635983), *Netta rufina* (USNM 292365), *Oxyura dominica* (USNM 430928), *Stictonetta naevosa* (USNM 612631), *Tadorna tadornoides* (USNM 638633), *Anser fabalis* (USNM 623342, USNM 643021), *Anser (Chen) caerulescens* (USNM 431549, USNM 345620), *Branta canadensis* (USNM 488584), *Cygnus atratus* (USNM 19738), *Coscoroba coscoroba* (USNM 346635) |
| **Extinct Taxa** | *Paakniwatavis grandei* (FMNH PA725), *Chaunoides* (Alvarenga 1999), *Diatryma* (*Diatryma giganteus* AMNH FR 70; Matthew and Granger 1917; Andors 1992), *Pelagornis chilensis* (Mayr and Rubilar-Rogers 2010), *Protodontopteryx ruthae* (Mayr et al. 2019), *Telmabates antiquus* (AMNH FR 3166-3186; Howard 1955), *Presbyornis* (*Presbyornis pervetus*: USNM PAL 483163-483166, USNM PAL 510082, USNM PAL 641330; *Presbyornis* sp.: USNM PAL ACC 2016973, USNM PAL ACC 335940, USNM PAL ACC 392324, USNM PAL ACC 393002, USNM PAL 498770-498771, USNM PAL 516605, USNM PAL 617185, USNM PAL 299845-299848, USNM PAL 618166-618180, USNM PAL 618183, USNM PAL 618189-618200, USNM PAL 618202, USNM PAL 618204-618207, USNM PAL 618209-618210, USNM PAL 618212-618215, USNM PAL 618218-618219, USNM PAL 618223-618224; *Presbyornis isoni*: USNM PAL 294117; Olson and Feduccia 1980; Elzanowski and Stidham 2010; CT scans from Zelenitsky et al. 2011 of *Presbyornis* sp. USNM PAL 200846), *Vegavis iaai* (Clarke et al. 2005, Clarke et al. 2016), *Conflicto antarcticus* (Tambussi et al. 2019), *Anatalavis oxfordi* (Olson 1999), *Gallinuloides wyomingensis* (Mayr and Weidig 2004), *Ichthyornis dispar* (Clarke 2004), *Lithornis promiscuus* (USNM PAL 336570, USNM PAL 424072, USNM PAL ACC 378571, USNM PAL 391983; Houde 1988; CT scans from Zelenitsky et al. 2011 of *Lithornis promiscuus* USNM PAL 391983), *Calciavis grandei* (Nesbitt and Clarke 2016), *Asteriornis maastrichtensis* (Field et al. 2020), *Wilaru tedfordi* and *Wilaru prideauxi* (De Pietri 2016) |

### Systematic Paleontology

AVES Linnaeus, 1758 *sensu* Gauthier and deQueiroz 2001 NEOGNATHAE Pycraft, 1900 *sensu* Gauthier and deQueiroz 2001 Anseriformes Wagler, 1831 *Paakniwatavis grandei*, gen. et sp. nov.

#### Holotype Specimen

FMNH PA725, a partial skeleton and tracheal rings preserved in a kerogen-poor laminated micrite slab (Figures 1 and 2). Measurements are provided in Table 2. Most of the vertebrae are absent or obscured where present. The shoulder girdle, thoracic vertebrae, ribs, pelvis, femora, synsacrum, caudal vertebrae and pygostyle have been eroded due to taphonomic processes. It appears that bacteria-induced or some other organic erosion of the bone has occurred. This type and extent of organic erosion is unique within recovered avian fossils from FBM. Similar taphonomy has been reported in an early Cretaceous Enantiornithine (Peteya et al., 2017) the early Cretaceous *Microraptor gui* (Hone et al., 2010), and several other Jurassic and Cretaceous avialan theropods (Currie and Chen, 2001; Hu et al., 2009), although much less erosion and deformation of the bone has occurred in these specimens compared to that of the holotype specimen of *P. grandei*. It has been suggested that this taphonomic phenomenon is due to changes in matrix chemistry caused by water being trapped between the feathers and the body (Hone et al., 2010). Scanning electron microscopy and additional analysis of the holotype specimen of *P. grandei* is necessary to determine the cause of this rare taphonomy.

**Figure 1.**
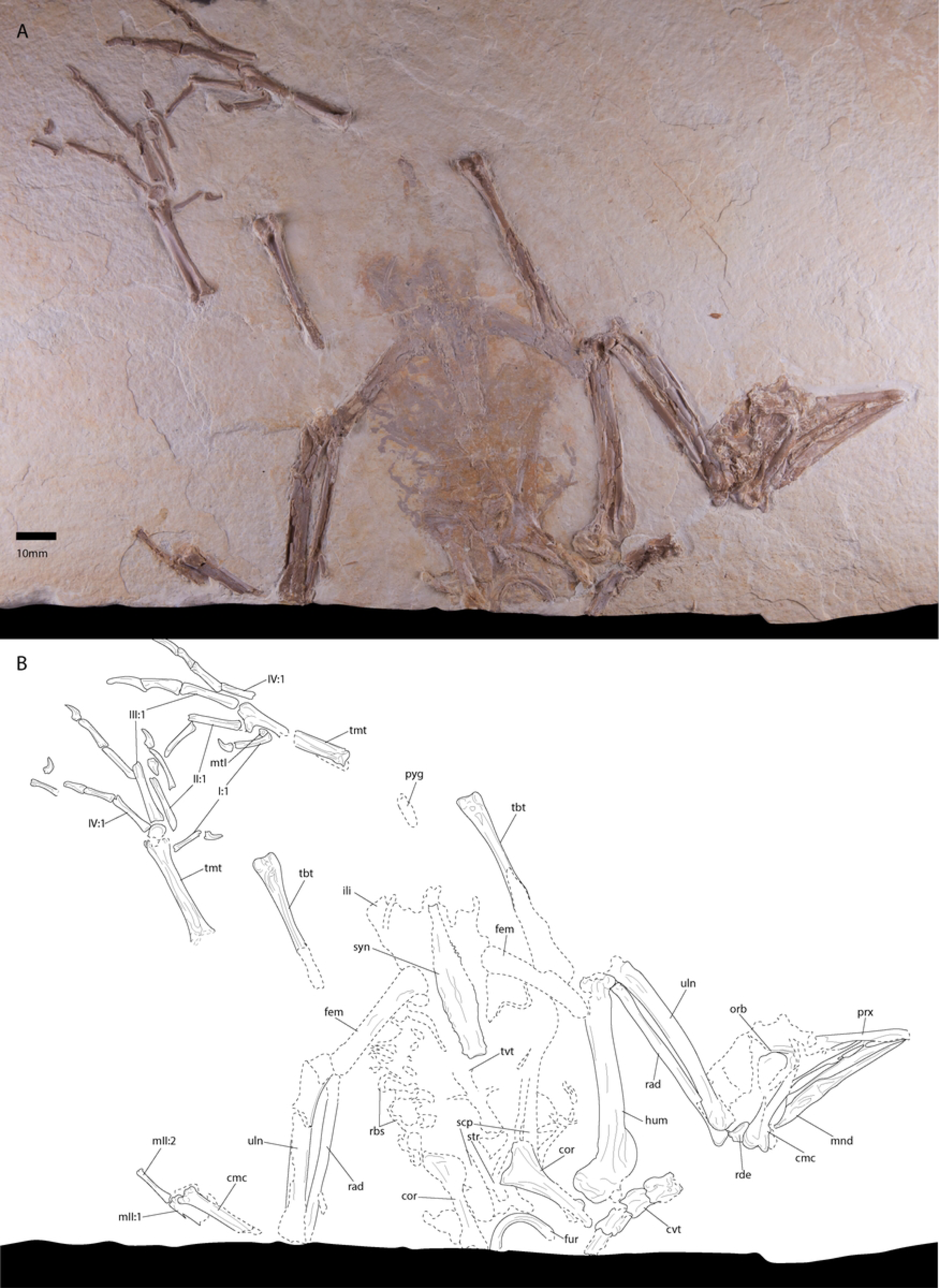
Photograph (A) and line drawing (B) of the holotype specimen of Paakniwatavis grandei (FMNH PA725). Extremely crushed bone and bone margin is delimited with dashed margins. Anatomical abbreviations: prx, premaxilla; orb, orbital margin; mnd, mandible; cvt, cervical vertebrae; tvt, thoracic vertebrae; syn, synsacrum; pyg, pygostyle; cor, coracoid; scp, scapula; fur, furcula; str, sternum; rbs, ribs; hum, humerus; uln, ulna; rad, radius; rde, radiale; cmc, carpometacarpus; mII:1, phalanx 1 of manual digit II; mtII:2, phalanx 2 of manual digit II; ili, ilium; fem, femur; tbt, tibiotarsus; tmt, tarsometatarsus; mtI, metatarsal I; I:1, phalanx 1 of pedal digit I; II:1, phalanx 1 of pedal digit II; III:1, phalanx 1 of pedal digit III; IV:1, phalanx 1 of pedal digit IV.

**Figure 2.**
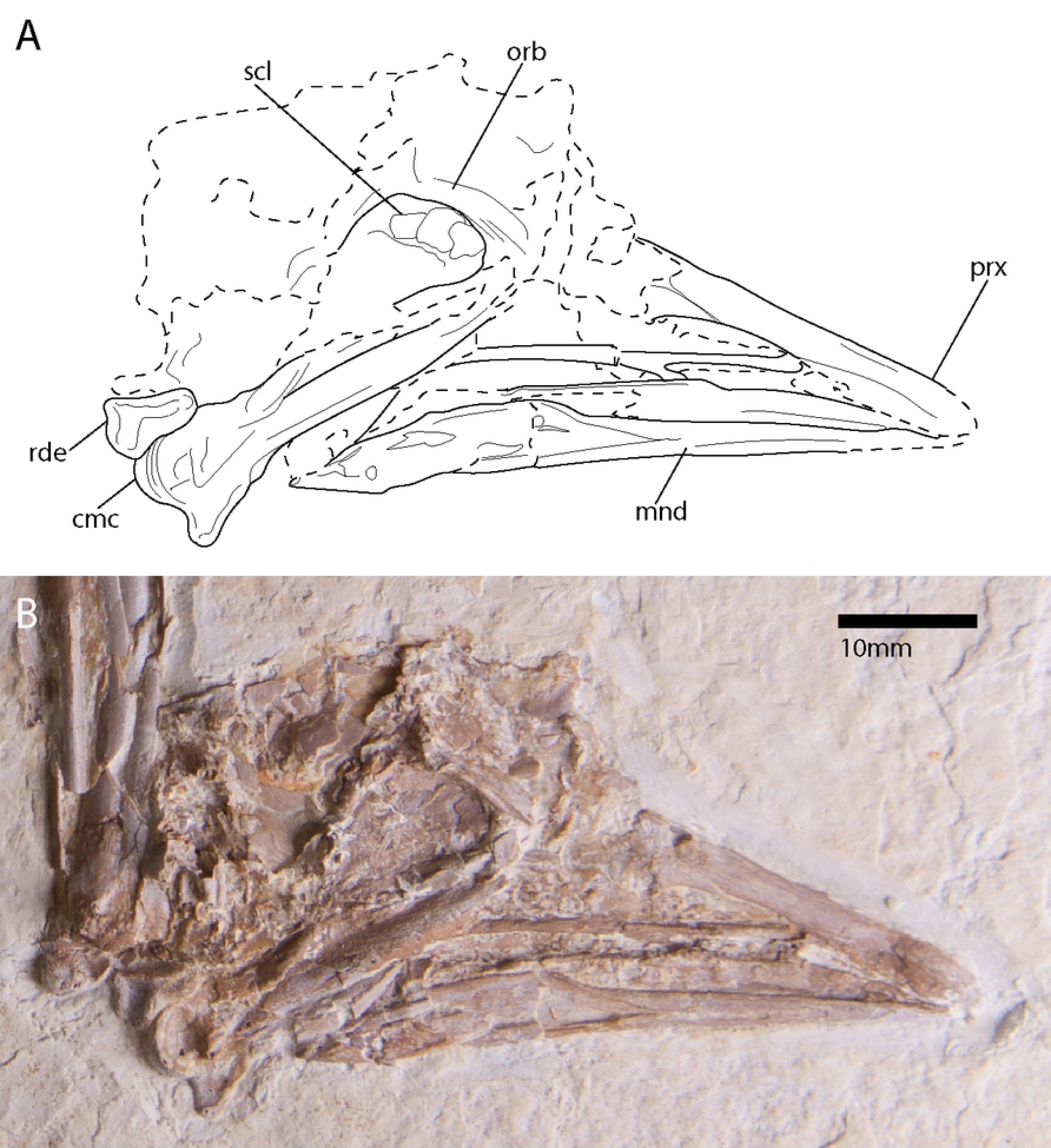
Photograph (A) and line drawing (B) of the holotype specimen of *Paakniwatavis grandei* (FMNH PA725). Bone is unfilled. Extremely crushed bone and bone margin is delimited with dashed margins. Anatomical abbreviations: prx, premaxilla; orb, orbital margin; scl, scleral ossicles; mnd, mandible; rde, radiale; cmc, carpometacarpus.

**Table 2.** Selected measurements of *Paakniwatavis grandei* in millimeters (mm), taken from surface of the holotype specimen slab (left/right) compared with taken and previously published measurements of *Presbyornis* (Olson and Feduccia 1980, Elzanowski and Stidham 2010), *Telmabates antiquus* (Howard 1955), and *Anatalavis oxfordi* (Olson 1999). Pedal phalanges are described using the format (digit:phalanx). Measurements are given for holotype specimens only.

| Measurement<br>(mm) | *Paakniwatavis*<br>*grandei* | *Presbyornis* | *Telmabates*<br>*antiquus* | *Anatalavis*<br>*oxfordi* |
| --- | --- | --- | --- | --- |
| Total skull length | 64.1 | 90 |  | 100.0 |
| Rostrum length | 28.2 | 38.4 |  |  |
| Furcula width | 19.3 |  |  | 7.0 |
| Coracoid length | 32.7/~29.6 | 33.5-35.3 | 43.8, 42.3 | 48.0 |
| Coracoid width at<br>midshaft | 5.44/4.0 |  |  | 8.7 |
| Humerus length | /71.3 |  |  | 119.3 |
| Humerus width<br>midshaft | /7.4 |  |  | 9.7 |
| Ulna length | 58.4/63.2 |  |  |  |
| Carpometacarpus<br>length | /~30.2 | 46.5-50 | 63.1 | 69.5 |
| Manual phalanx II:<br>digit 1 | 15.5/ | 19.4 | 29.6, 30.4 | 30.2 |
| Manual phalanx II:<br>digit 2 | 9.6/ |  |  | 22.7 |
| Synsacral length | 50.3 |  |  | >52 |
| Pelvis acetabular width | 19.5 | 5.1-6.4 |  |  |
| Femur length | 40.6/41.0 | 57-62 | 69-72 |  |
| Tibiotarsus length | /66.1 | 120<br>estimated |  |  |
| Tarsometatarsus length | 38.6/38.1 | 105-120<br>estimated<br>(minimum 93) | Minimum 105<br>estimated |  |
| I:1 length | 9.5/11.1 |  |  |  |
| II:1 length | 7.4/6.7 |  |  |  |
| II:2 length | 13.4/14.3 |  |  |  |
| III:1 length | 19.8/18.5 |  |  |  |
| III:2 length | 13.7/13.1 |  |  |  |
| III:3 length | 11.7/9.7 |  |  |  |
| IV:1 length | 14.2/11.2 |  |  |  |
| IV:2 length | 9.1/10.2 |  |  |  |
| IV:3 length | 6.9/7.8 |  |  |  |
| IV:4 length | 8.8/7.3 |  |  |  |

The holotype specimen was scanned using dual tube x-ray computed tomography at the PaleoCT Lab at the University of Chicago, which can scan specimens with a resolution of up to 0.4 μm. As the specimen slab was large, it was scanned using a two-part multiscan that was combined to form one image sequence. The voxel size of the combined scan is 103.8710. The specimen is housed in the Department of Geology of FMNH. CT data generated during the current study are available in the Supplementary Data via Morphobank (O’Leary and Kaufman, 2012) under Project 4001 (http://morphobank.org/permalink/?P4001).

#### Etymology

*Paakniwatavis* references *Paakniwat*, used by the Shoshoni tribe indigenous to the region of the recovery site and means “Water Spirit” (Shoshoni Language Project, 2018). The Water Spirits are dangerous supernatural beings that lure people to their death with child-like cries. The name references the aquatic ecology of this taxon. The species honors Dr. Lance Grande, who collected the holotype specimen, in recognition of his leading research on the faunas of the Green River Formation.

#### Type locality and horizon

The holotype specimen was collected from FBM (*sensu* Buchheim, 1994) Locality H (F-2 H in Grande and Buchheim, 1994; Grande, 2013). FBM Locality H is one of several near-shore localities that have produced avian fossils, and is located in the northeastern near-shore region. Locality H is within a four meter thick horizon representing a few hundred to a few thousand years of the early Eocene (Grande and Buchheim, 1994). The horizons of the near-shore deposits of FBM are thicker than those of the mid-lake localities due to increased sedimentation near the shore. The fossil-bearing KPLM facies are characterized by thick kerogen-poor calcite laminae and instances of thin organic laminae. Laminae alterations can be differentiated by inconsistent texture where the organic laminae is absent (Grande and Buchheim, 1994). Locality H has thus far yielded the highest number of avian fossils comprising lithornithids (palaeognathid; Nesbitt and Clarke, 2016), *Gallinuloides wyomingensis* Eastman 1900 (see also Weidig, 2010; galliform), two frogmouth-like specimens (Nesbitt et al., 2011), a possible oilbird (Olson, 1987), four frigate birds (Olson, 1977; Olson and Matsuoka, 2005; Stidham, 2015), an ibis (Smith et al., 2013), a turaco (Field and Hsiang, 2018), two *Messelornis nearctica* specimens (Hesse, 1992; Weidig, 2010), a jacamar-like bird (Weidig, 2010), a hoopoe- like bird (Grande, 2013), stem rollers (Coraciiformes; Ksepka and Clarke, 2010), stem parrots (Ksepka and Clarke, 2011), additional taxa within Telluraves (Feduccia and Martin, 1976; Ksepka et al., 2019), and several unknown birds (Grande, 2013). The near-shore deposits are additionally characterized by juvenile fish being more commonly preserved, abundant benthonic invertebrates, stingrays (Batoidea), lizards, crocodiles, turtles, and non-flying mammals. Locality H also has the only known amphibian preserved within FBM (Rieppel and Grande, 1998).

#### Diagnosis

*Paakniwatavis grandei* is diagnosed by a proposed unique combination of characters comprising (1) a mediolaterally narrow rostrum (character 3:state 1; Figures 1-3), (2) an elongate and dorsoventrally thick retroarticular process (Figure 3; 241:2, 254:2), (3) a dorsoventrally thick furcula (Figure 3; 424:2), (4) thoracic vertebrae that are not solely heterocoelous (286:2), (5) presence of a supracoracoid nerve foramen (Figure 4; 391:1), (6) lack of a spur on the carpometacarpus (509:1), (7) femora that are half the length of the tibiotarsi (Figure 1; 597:1), (8) presence of a prominent tubercle laterodistal to the pons supratendinous of the tibiotarsus (643:1), (9) tarsometatarsi that are just over half the length of the tibiotarsi (Figure 1; 656:1), (10) a medial hypotarsal crest that is projected farther plantar than the lateral crest (Figure 4; 668:1), and (11) a deep sulcus extensorius of the tarsometatarsus (Figure 1; 686:2). Diagnosis for the genus as per the species.

**Figure 3.**
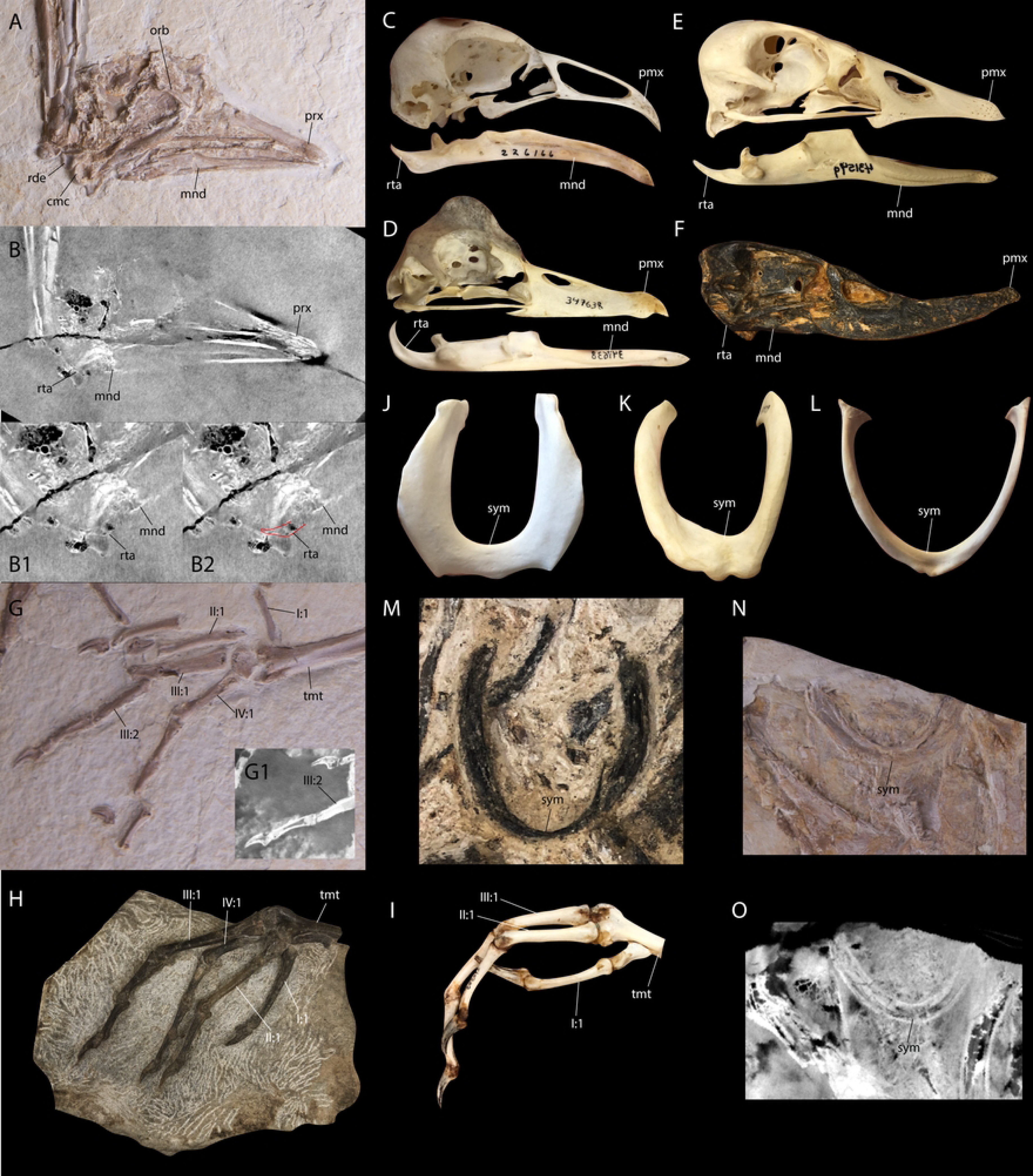
Comparison of anseriform traits in *Paakniwatavis grandei* (FMNH PA725) to those of the extinct *Presbyornis* and extant Anseriformes (*Chauna torquata*, *Anseranas semipalmata*, and *Chloephaga melanoptera*). Photograph (A) and CT scan slice (B) of the holotype specimen of *P. grandei* (FMNH PA725), with close-ups (B1 and B2) of CT scan slices of the caudal mandible. Photographs of the skull and mandible of (C) *C. torquata*, (D) *A. semipalmata*, (E) *C. melanoptera*, and (F) *Presbyornis sp.* (USNM 299846). Photograph (G) and CT scan slice (G1) of the left tarsometatarsus and pedal phalanges of *P. grandei*. Photographs of (H) the left tarsometatarsus and pedal phalanges of *Presbyornis sp.* (USNM ACC 335940) and (I) the right tarsometatarsus and pedal phalanges of *C. torquata.* Photographs of the furcula of (J) *C. torquata*, (K) *A. semipalmata*, (L) *Anas platyrhynchos*, and (M) *Presbyornis sp.* (USNM ACC335940). Photograph (N) and CT scan slice (O) of the furcula of *P. grandei* from the holotype specimen. Anatomical abbreviations: prx, premaxilla; orb, orbital margin; mnd, mandible; rta, retroarticular process; rde, radiale; cmc, carpometacarpus; sym, symphysis; tmt, tarsometatarsus; I:1, phalanx 1 of pedal digit I; II:1, phalanx 1 of pedal digit II; III:1, phalanx 1 of pedal digit III; III:2, phalanx 2 of pedal digit III; IV:1, phalanx 1 of pedal digit IV.

**Figure 4.**
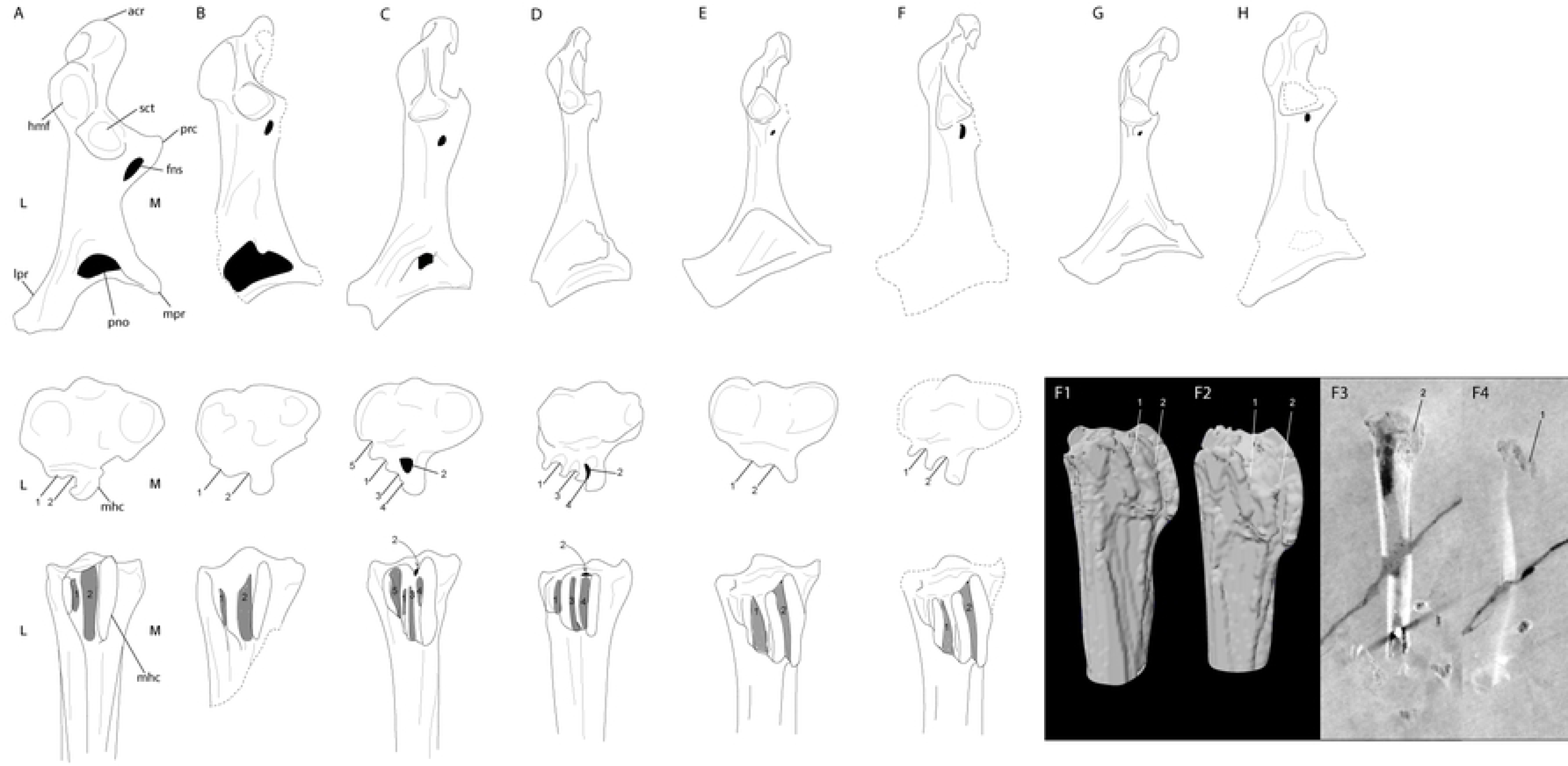
Comparison of the coracoid and hypotarsus of the tarsometatarsus of *Paakniwatavis grandei* (FMNH PA725) to those of the extinct *Presbyornis*, *Telmabates antiquus*, *Chaunoides antiquus* and *Anatalavis oxfordi* and extant Anseriformes. Line drawings of the coracoids and hypotarsi of (A) *Anhima cornuta*, (B) *C. antiquus*, (C) *Anseranas semipalmata*, (D) *Dendrocygna guttata*, (E) *Presbyornis sp.,* (F) *P. grandei*, (G) *T. antiquus* and (H) *A. oxfordi*. Coracoids are depicted in dorsal aspect, hypotarsi are depicted in proximal and plantar aspects. L and M in panel (A) denote the lateral and medial sides of each element. (F1 and F2) are segmented hypotarsi of *P. grandei* in lateroplantar aspect. (F3 and F4) are CT scan slices of the same tarsometatarsus of *P. grandei*. Anatomical abbreviations: acr, processus acrocoracoideus; hmf, humeral facet; sct, scapular facet; prc, procoracoid process; fns, supracoracoid nerve foramen; pno, pneumatic opening; lpr, lateral process; mpr, medial process; mhc, medial hypotarsal crest; 1, sulcus for tendon of musculus flexor hallucis longus (fhl); 2, sulcus or canal for tendon of musculus flexor digitorum longus (fdl); 3, sulcus for tendon of musculus flexor perforatus digiti 2 (fp2); 4, sulcus for tendon of musculus flexor perforans et perforatus digiti 2 (fpp2); 5, sulcus for muscularus fibularis longus (fbl).

#### Differential Diagnosis

The rostrum shape exhibited by this taxon is unlike any other previously recovered Paleogene Anseriformes. *Paakniwatavis* is easily distinguished from *Anatalavis* and *Presbyornis* by this feature. Unlike the mediolaterally wide, duck-like bills of *Anatalavis* and *Presbyornis* (mediolaterally wider than the width across the paroccipital processes), *Paakniwatavis grandei* exhibits a mediolaterally narrow bill that is narrower than the width of the skull at the parocciptal processes and, in this feature, is more like that of Anhimidae (Figures 1-3). The tomial margins of the bills of *Presbyornis* and *Anatalavis* are similarly dorsoventrally thick and recurved, unlike the straight and dorsoventrally narrow facial margin of the bill in *Paakniwatavis* (Figure 3). The nares of *Paakniwatavis* are over half the length of the rostrum, whereas those of *Presbyornis* and *Anatalavis* are less than half of their rostral length. The synsacral count of *Paakniwatavis* is within 14-19 vertebrae, whereas it is within 10-13 for *Telmabates* and *Presbyornis*. In the coracoid, a small blind pneumatic foramen directly below the scapular cotyla is present in *Presbyornis* and *Telmabates* but is absent in *Paakniwatavis* and *Anatalavis*. The coracoid in *Paakniwatavis* is more elongate relative to width of the sternal facet than that of *Anatalavis* (Figure 4). The rami of the furcula are extremely thick in *Paakniwatavis* compared to the thin furculae of *Presbyornis* and *Anatalavis* (Figure 3). The acromion process of the scapula is truncate, unlike the cranially elongate processes of *Telmabates* and *Presbyornis*.

The dorsal angle of the scapula is caudal to the midpoint of the shaft in *Telmabates* and *Presbyornis*, but is at the midpoint in *Paakniwatavis.* The incisura capitis of the humerus is deep in *Paakniwatavis* and *Presbyornis* but shallow in *Telmabates.* The fossa pneumotricipitalis of the humerus is pneumatic in *Paakniwatavis* and *Anatalavis* but apneumatic in *Telmabates* and *Presbyornis*. *Paakniwatavis* has a deeper impression coracobrachialis cranialis than *Telmabates* or *Presbyornis*. The sulcus ligamentosus transversus is more truncate in *Paakniwatavis* than in *Presbyornis* or *Anatalavis*. The bicipital crest is shorter than half the length of the deltopectoral crest in *Paakniwatavis*, whereas it is over half this length in *Presbyornis* and *Telmabates*. The fossa olecrani of *Paakniwatavis* is more shallow than those of *Presbyornis* and *Anatalavis*.

*Paakniwatavis* has a deeper fossa infratrochlearis of the carpometacarpus than *Anatalavis*. The craniocaudal lengths of manual digits II and III at the synostosis are subequal in *Paakniwatavis*, whereas II is greater than III in *Anatalavis.* The epicondylus medialis of the tibiotarsus is more pronounced in *Paakniwatavis* than in *Telmabates* or *Presbyornis.* The epicondylus medialis is less pronounced than those of *Presbyornis* or *Chaunoides antiquus* Alvarenga 1999.

*Paakniwatavis* can be differentiated further from *Presbyornis* based on features of the quadrate, mandible, humerus, ulna, carpometacarpus, tibiotarsus, tarsometatarsus, and pedal phalanges. The crista tympanica of the quadrate terminates within the ventral half of the quadrate in *Paakniwatavis*, whereas it terminates within the dorsal half in *Presbyornis*. The tuberculum subcapitulare is separated from the squamosal capitulum in *Paakniwatavis*, but is contiguous with the capitulum in *Presbyornis*. The relative heights of the rostral and caudal apices of the coronoid process of the mandible are subequal in *Paakniwatavis*, but the rostral apex is higher in *Presbyornis*. The mandibular ramus caudal to the rostral fenestra mandibulae is deep and concave along the medial face in *Paakniwatavis*, and is relatively shallow in *Presbyornis*. A mandibular ventral angle is prominent in *Prebyornis*, but not in *Paakniwatavis.* While the fenestra rostralis mandibulae is slit-like in *Paakniwatavis*, it is transverse and largely perforate in *Presbyornis* (Figure 3). The mandibular process in *Paakniwatavis* is exceptionally tapered, unlike the robust process of *Presbyornis*. *Paakniwatavis* has narrower crista deltopectoralis of the humerus than *Presbyornis*. The fossa m. brachialis is located more medially in *Paakniwatavis*. The depression radialis of the ulna is deeper in *Paakniwatavis.* The labrum dorsalis is of the carpometacarpus is more sharply angled in *Paakniwatavis.* The pons supratendinous of the tibiotarsus opens along the midline in *Paakniwatavis* rather than medially. The tarsometatarsus is approximately half the length of the tibiotarsus or less in *Paakniwatavis*, whereas the length of these elements is subequal in *Presbyornis*. The lateral cotyle of the tarsometatarsus is more shallow in *Paakniwatavis.* In *Paakniwatavis*, pedal phalanx IV: digit IV is longer than IV: III, whereas the opposite is true in *Presbyornis*.

*Paakniwatavis* can be differentiated further from *Telmabates* based on features of the humerus and carpometacarpus. *Paakniwatavis* has a domed crista along the proximal margin of the fossa pneumotricipitalis of the humerus, whereas this crista is typical in *Telmabates*. The impression m. pectoralis is deeper in *Paakniwatavis*. The trochlea carpalis of the carpometacarpus is deeper in *Paakniwatavis.* The epicondylaris medialis depression is more shallow in *Paakniwatavis* than in *Telmabates*.

*Paakniwatavis* can be differentiated form Chaunoides due to differences in the coracoid and tarsometatarsus. Within the coracoid, the primary axis of the scapular cotyla is skewed laterally in *Paakniwatavis*, *Presbyornis*, *Telmabates* and *Anatalavis* but is centralized in *Chaunoides*. The sternal facet curves cranially in *Chaunoides* but is flat in *Paakniwatavis*. In the tarsometatarsus, the major hypotarsal ridge in *Paakniwatavis* is hooked distally.

### Description and Comparison

#### Skull and Mandible

The skull and rostrum are preserved in dorso-lateral aspect. The right carpometacarpus has broken through the skull and mandible and caused deformation along the caudoventral margins of these elements. The bill length is roughly equal to that of the cranium. It is mediolaterally narrow and tapered toward the anterior margin as in Anhimidae. This is unlike the mediolaterally broad, duck-like bills of most anseriform-like fossils such as *Anatalavis* and *Presbyornis*. The terminus of the rostrum is slightly decurved. The caudal and anterior portions of the nares are largely broken, but CT scans (see Supplementary Data) show that the nares would have been holorhinal and rostral to the zona flexoria craniofacialis (ZFC) as in all Galloanserines. The length of the nares is over half that of the rostrum, a condition only present in Anseriformes within Anhimidae. The cranium immediately caudal to the ZFC appears to have had the pneumatized swelling seen in *Chauna* which is described in Livezey (1997, character 10). The overall shape and size of the cranium is most similar to those of Anhimidae or Anseranatidae. The robust left jugal is present between the rostrum and left ramus of the mandible. A prominent, lateroventrally projecting supraorbital crest is present, like those of Anhimidae (see Musser and Cracraft 2019). CT data reveals that the postorbital process is elongate as in most Anseriformes (see Supplementary Data). The zygomatic process is absent as in all Anseriformes. At least three large, broad scleral ossicles have been preserved along the rostral margin of the orbit. Fonticuli within the interorbital area appear to be absent.

The mandible is dorsoventrally thin along the proximal margin and widens caudal to the coronoid process. While both rami are visible on the surface of the slab, much of the left mandible and the left quadrate are obscured by the carpometacarpus and are only visible within the CT data (see Supplementary Data). The rostral terminus of the mandible is broken and it cannot be assessed whether it was decurved as in Anhimidae. The rostral mandibular fenestrae are rostrocaudally elongate and slit-like but appear to open rostrally. The coronoid processes are tuberculate and subtle compared to those of most Anseriformes. CT data reveals an elongate medial process at the condylar area of the mandible as in all Galloanserines. On the right ramus, a slender, recurved retroarticular process that has been severed from the mandible by the carpometacarpus can also be seen in the CT data (see Supplementary Data). A retroarticular process is present in all Galloanserines and included fossils.

The left quadrate is visible in the CT data (see Supplementary Data). Most features of the quadrate have been obliterated, including the orbital crest. No foramen is present between the capitulae, but a foramen is present on the medial face of the otic process as in all Anseriformes. A tuberculum subcapitulare is present as in all Galloanserines. A prominentia submeatica appears to be present as in all Anseriformes.

#### Axial Skeleton

The atlas, axis and the third cervical vertebra are poorly preserved in ventrolateral aspect near the furcula, with two additional cervical vertebrae preserved caudal to the third cervical vertebra in ventral aspect. The dorsal spines are rounded and not dorsoventrally prominent. This condition is more similar to that of Anseranatidae rather than that of Anhimidae. The third cervical vertebra appears to be more elongate and to have a less prominent dorsal spine, suggesting that the cervical series elongates caudally.

The thoracic and pelvic areas of the specimen are poorly preserved. The bone appears to have eroded due to taphonomic processes, possibly due to bacterial erosion. Several caudalmost cervical vertebrae and thoracic vertebrae are visible on the surface of the slab and within the CT data (see Supplementary Data). Most of the thoracic vertebrae are preserved in ventral aspect.

CT data reveals that the thoracic series is not completely heterocoelous. This condition is present in *Presbyornis*, *Anatalavis* and *Telmabates* but lost in extant Anseriformes. The thoracic vertebrae do not fuse to form a notarium. This condition is present in all extant Anseriformes with the exception of Anseranatidae.

The synsacrum is preserved in ventral aspect. CT data reveals that at least 14 synsacral vertebrae are present (see Supplementary Data). This is the condition in all extant Anseriformes. *Telmabates* and *Presbyornis* have 13 or fewer synsacral vertebrae. The sulcus ventralis of the synsacrum is present and appears to have been deep. Several poorly preserved caudal vertebrae with indiscernible features are present in the CT data (see Supplementary Data). A stout pygostyle is visible caudal to the synsacrum on the surface of the slab.

#### Shoulder girdle

The symphysis and ventral clavicles of the furcula are preserved in caudal aspect. The furcula is more robust than those of most Anatidae but more gracile and thin than those of Anhimidae or Anseranatidae (Figure 3). A processus interclavicularis dorsalis is absent, and an apophysis is absent.

Both coracoids are preserved in ventral aspect. The distal bases and shafts of the coracoids can be seen. The acrocoracoid process is robust and hooked. CT data reveals a procoracoid process and supracoracoid nerve foramen to be present. Portions of the scapulae are present near to or overlapping the coracoids, but most of their morphology is indiscernible.

#### Forelimbs

The right humerus is preserved in cranio-medial aspect. The sulcus ligamentosus transversus is extremely deep. The crista deltopectoralis is prominent, rounded and flares cranio-laterally like those of Anseriformes. The humeral head is bulbous and prominent. Also as in *Anas*, the impression of the coracobrachialis is relatively deep. The condylus dorsalis is small and hamate. The tuberculum supracondylare ventral is large and bulbous like that of *Anas.* The epicondylus dorsalis appears to be distally extensive, and most similar to that of *Anas*. CT data reveals the morphology of the caudal humerus; the crista proximally edging the fossa pneumotricipitalis appears to have been domed slightly, like that of *Anatalavis*.

The right radius and ulna are preserved in ventral aspect. Most features of these bones cannot be seen or were not preserved. The ulna body is thick and robust, but shorter than the humerus.

The olecranon is much shorter and more rounded than that of *Anas*, and is most similar in size and shape to that of *Chauna.* The impression brachialis appears to be proximally deep and ovoid, similar those of Anhimidae. Much of the distal portion of the ulna is obscured by the skull. There appears to be small pneumatic foramen just under the cotyla humeralis of the radius. The left ulna and radius are present but not as well preserved as those of the right. They are preserved in dorsal aspect.

The right carpometacarpus overlaps the mandible and has broken onto the skull. It is preserved in ventral aspect. The processus pisiformis is elongate, rounded and caudally oriented; it is very similar to that of *Anseranas.* The rim of the dorsal trochlea is prominent and strongly angled, which is also similar to the condition of *Anseranas*. The processus extensorius is identical in size and shape to that of *Anseranas* as it is triangular in shape with a rounded point. The os carpi ulnare is present as well and is visible in the CT scans (see Supplementary Data).

The os metacarpale minus is obscured by the skull but appears to be dorso-ventrally thick. Only the distal shaft and condylar area of the left carpometacarpus is preserved and is in cranial aspect. The facies articularis digitalis major on the left carpometacarpus form a 3-pronged distal end with rounded termini. A prominent crista is present towards the dorsal aspect of the digital articular area; it is not seen in *Anas* and in *Chauna* it is hook-like. The os metacarpale minus appears broken and only a small spatium intermetacarpale appears present, but this could be exaggerated by crushing. The phalanges digitus majoris are preserved in dorsal aspect, although digit I is somewhat obscured by the carpometacarpus. The pila cranialis of phalanx dig. majoris I appears robust. Phalanx II of this digit is thin and elongate.

#### Sternum

The sternum is extremely poorly preserved but appears to have been broad and subrectangular like that of *Anas*. A prominent carina is distorted but preserved. An elongate spina externa appears to be present, but it cannot be discerned with confidence whether this is part of the sternum or a vertebra lying under the sternum. What appears to be an isolated uncinate process looks visible on a rib located to the left of the sternum; however, this also cannot be assessed with confidence.

#### Pelvic girdle

The pelvis is preserved in ventral aspect. A pair of acetabular struts are visible in the CT data. Portions of the postacetabular ilium, ischium and a medially curved pubis can be seen more clearly in the CT data.

#### Hindlimbs

The right and left femora are poorly preserved. The head of the right femur appears robust like those of Anhimidae or Anseranatidae. The femora are half the length of the tibiotarsi. The right and left tibioarsi are preserved in cranial view. All that remains of the left tibiotarsus is the mid and distal shaft and the condylar area. The bodies of the tibiotarsi are long and slender. The entire right tibiotarsus appears is preserved along with the head and proximal shaft of the right fibula. The cnemial crest is obscured by the right femur, but appears prominent and acuminate, very similar to that of *Anseranas*. The canalis and sulcus extensorius appear to be deep, and the pons supratendinus is ossified. No intratendinous ossification is present.

The tarsometatarsi are just over half the length of the tibiotarsi and are preserved in medio- dorsal aspect. The hypotarsus is visible in the CT data. The medial hypotarsal crest is more prominent, as in most Anseriformes. Two sulci are present. Both tarsometatarsi exhibit deep extensor sulci that extend to the distal portion of the tibiotarsus. In the right tarsometatarsus, metatarsal trochlea III reaches the most distally. Metatarsal trochlea II is deflected plantarly, as in Anatidae. The phalanges are elongate like those of *Anas*. IV is shorter than III but the hallux is elongate. Phalanx III is longer than the length of the tarsometatarsus.

#### Body mass

Femur length was the best predictor of body mass in extant volant birds (R2 = 0.9028; Field et al., 2013) that could be obtained from FMNH PA725, *Presbyornis* (Olson and Feduccia, 1980; Elzanowski and Stidham, 2010) and *Telmabates* (Howard, 1955). Average femur length measurements were used to calculate body mass for each taxon. An approximate mean body mass estimate for *P. grandei* is 304.4g based on the published allometric equation using femoral length from Field et al., 2013. The approximate mean body mass estimates for *Presbyornis* and *Telmabates* are 882.2g and 1423.0g, respectively based on the same equation.

### Phylogenetic Analyses and Ancestral State Estimation

#### Comparative Materials

Specimens used for the description and phylogenetic analyses came from the Bird Division of FMNH, the Ornithology Department of AMNH, the Ornithology Department of USNM, and the Texas Memorial Museum. Osteological terminology largely follows Baumel and Witmer (1993). Specimen numbers for examined taxa are presented in Table 1. Extinct taxa were scored from direct observation where possible. CT scans of the skulls of *Presbyornis* and *Lithornis promiscuus* were loaned from USNM for examination (originally produced for Zelenitsky et al., 2011). All USNM *Presbyornis* material and all AMNH *Telmabates antiquus* material was directly examined. Extinct taxa scored from photographs when necessary comprise *Ichthyornis dispar* Marsh 1872 (Clarke 2004), *Diatryma gigantea* and *Diatryma steini* (Matthew and Granger 1917; Andors 1992) *Pelagornis chilensis* (Mayr and Rubilar-Rogers 2010), *Protodontopteryx ruthae* (Mayr et al. 2021), *Vegavis iaai* (Clarke et al. 2005, Clarke et al. 2016), *Conflicto antarcticus* (Tambussi et al. 2019), *Anatalavis oxfordi* (Olson 1999), *Wilaru tedfordi* Boles 2013 and *Wilaru prideauxi* (De Pietri 2016), *Presbyornis* (Olson and Feduccia 1980, Elzanowski and Stidham 2010), *Chaunoides antiquus* (Alvarenga 1999), *Lithornis promiscuus* (Houde 1988), *Calciavis grandei* (Nesbitt and Clarke 2016), *Asteriornis maastrichtensis* (Field et al. 2020), and *Gallinuloides wyomingensis* (Mayr and Weidig 2004). Scorings for separate *Diatryma*, *Presbyornis* and *Wilaru* species were concatenated into genus-level taxa for more robust phylogenetic placement, especially since most recovered *Presbyornis* specimens have not been assigned to the species level.

#### Character Matrices and Ancestral State Reconstruction

The morphological data matrix is built on that of Musser and Cracraft (2019) and Musser and Clarke (2020) following the methodology discussed in those publications, but was modified to comprise 719 discrete characters and 41 taxa, 16 of which are extinct. Most of the characters detail skeletal morphology, although several characters additionally describe musculature, the syrinx, behavior, and ecology. Ten additional characters were added to the morphological matrix for ancestral state reconstruction of relevant traits that detail habitat preference, swimming mode, diet, status of rhamphothecal lamellae, status of pedal webbing, feeding mode, syrinx anatomy, and the relative length of pedal phalanx III compared to that of the tarsometatarsus. Ancestral state reconstructions were performed in Mesquite (The Mesquite Team 2021) using parsimony methods. All character descriptions are provided in the Appendix and all data matrices and analyses logs have been made publicly available on Morphobank (O’Leary and Kaufman 2012) under Project 4001 (http://morphobank.org/permalink/?P4001).

Characters from several previously published large-scale morphological datasets focused on early avian divergences and galloanserine-like fossils (Livezey, 1997; Ericson, 1997; Cracraft and Clarke, 2001; Clarke and Norell, 2001; Mayr and Clarke, 2003; Livezey and Zusi, 2006; Elzanowski and Stidham, 2010; Worthy et al., 2017) were evaluated for use in this iteration of the dataset, and characters from these matrices have been cited where a character was incorporated from a previously published matrix or where characters overlap with our matrix.

Characters were additionally used from several previously published matrices which have also been cited in the Appendix.

We additionally created a combined data matrix comprising the morphological data coupled with all available mitochondrial genomes available on GenBank (Clark et al., 2016; Sun et al. 2017) for this taxon sampling and the Early Bird II dataset from Reddy et al. (2017). The combined dataset has a total of 158,368 base pairs. This data matrix is also publicly available under the same project on Morphobank (O’Leary and Kaufman 2012). Sequences included were matched to the species level where possible; otherwise sequences of taxa within the same genus or family level were used. Mitochondrial genomes were aligned using MAAFT (Katoh et al., 2002). The mitochondrial genome data was not partitioned. Jmodeltest (Posada, 2008) was used to find the best fit substitution model, GTR+I+G as is common in vertebrate nonpartitioned mitochondrial genome data (e.g. Song et al., 2016). We used the GTR + G model for the Early Bird II dataset as in Reddy et al. (2017) but did not partition the dataset due to using a limited taxon sample.

#### Phylogenetic Analyses

We performed unconstrained heuristic parsimony analyses of the morphological dataset in PAUP* (Swofford, 2002) Version 4.0a, build 169 using 10,000 random taxon addition replicates per run. A backbone constraint that minimally constrained Galloanseres to be monophyletic needed to be employed when *Wilaru* was included, as inclusion of this taxon placed Galliformes as the sister taxon of included Palaeognathae. Heuristic search algorithms were used. Tree bisection reconnection branch swapping was employed and minimum branch lengths valued at zero were collapsed, following Mayr and Clarke (2003) and Musser and Cracraft (2019). No character weighting was applied. All characters were unordered. Bootstrap analyses were performed using 500 bootstrap replicates each with 10 random taxon addition replicates as in Mayr and Clarke (2003).

Within the combined data analysis, mitochondrial genomes were analyzed using a GTR + invariable gamma substitution model (Lanave et al., 1984; Yang, 1994). The Early Bird II dataset was analyzed using a GTR + gamma substitution model. Partitions and model settings are detailed in the available matrix files. The Mk model (Lewis, 2001) was used for the morphological data partition within our combined data matrix. Bayesian analyses (Yang and Rannala, 1997) of molecular and combined data were performed in MrBayes (Huelsenbeck and Ronquist, 2001; Ronquist and Huelsenbeck, 2003; Version 3.2.7a) via the CIPRES portal (Miller, Pfeiffer and Schwartz, 2010). MrBayes settings used were default, with the exception of running the analysis for 40,500,000 generations. The Bayesian analysis code as well as all matrices and output files are included in the Supplementary Data.

### Principle Component and Linear Discriminant Analyses

Principle component analysis (PCA) of skeletal measurements of relevant taxa from Hinic-Frlog and Motani (2010) was conducted in R (RStudio Team 2020) to assess swimming mode based on skeletal measurements. We added *P. grandei*, *Chauna torquata*, *Telmabates antiquus*, *Presbyornis*, *Anatalavis oxfordi*, and *Vegavis iaai* to this dataset. Measurements for the latter four taxa were largely taken from the literature (see Table 1). Linear discriminant analysis was additionally performed on the same data in R. R code and measurement data are available in the Supplementary Data.

## Results

*Paakniwatavis grandei* is recovered across all analyses as the sister taxon of Anseranatidae (∼50% bootstrap value in both morphological results, 85% clade credibility in both combined results; see Figure 5 and Supplementary Figures on Morphobank). Across both the parsimony analysis of morphological data and the Bayesian analysis of combined data, addition of *Wilaru* results in differing topologies (see Supplementary Data). Within the morphological results, addition of *Wilaru* results in its placement as the sister taxon of *Diatryma* within a stem anseriform group (<50% bootstrap support) and placement of *Asteriornis maastrichtensis* Field et al. 2020 as the sister taxon of a clade containing Tinamiformes+Lithornithidae (100% bootstrap support) rather than the sister taxon to Galloanseres (<50% bootstrap support) as in the morphological results that excluded *Wilaru*.

**Figure 5.**
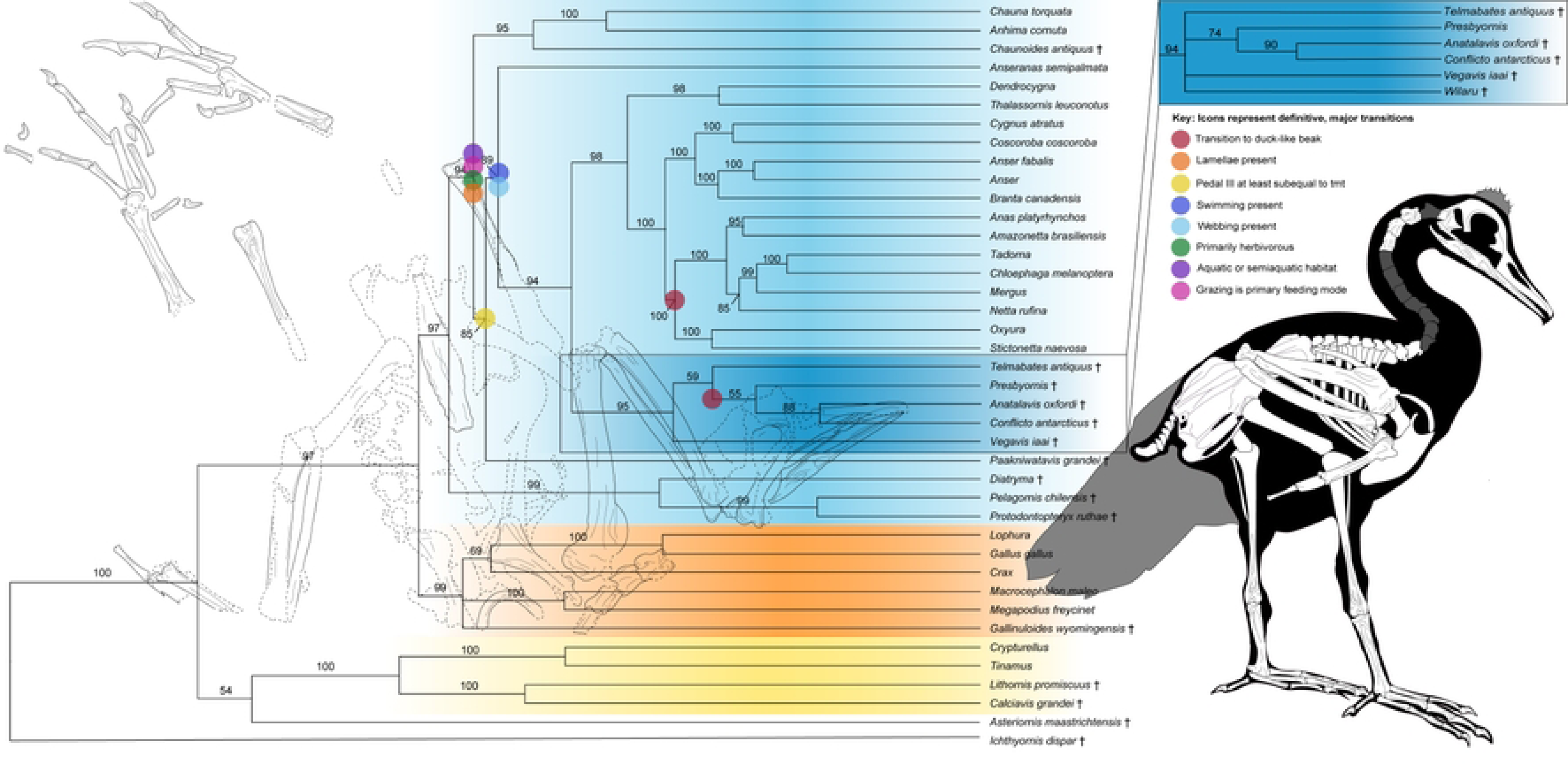
Resulting consensus tree from Bayesian analysis of 719 morphological characters and 158,368 base pairs. Clade credibility values greater than 50% are annotated above branches. Extinct taxa are delimited with daggers. A line drawing of the holotype specimen of *Paakniwatavis grandei* is overlaid on the tree and a reconstruction of this species is shown to the right of the tree. Icons represent definitive ancestral state reconstruction of the earliest transitions. Only the first major transitions for Anseriformes are shown, with subsequent transitions excluded. The inset to the right of the tree displays how results changed when *Wilaru* was included in analysis.

Due to this, the monophyly of Galloanseres was minimally constrained using a backbone constraint when *Wilaru* was added to the morphological analysis, and the following results discussed with *Wilaru* included comprise a monophyletic Galloanseres. Addition of *Wilaru* changes the relationships within the stem anatid clade containing *Vegavis iaai*, *Telmabates*, *Presbyornis*, *Conflicto*, and *Anatalavis oxfordi* recovered across all analyses. With *Wilaru* excluded, *Telmabates* and *Presbyornis* are placed as sister taxa and form the sister clade to a group comprising *Vegavis*(*Conflicto+Anatalavis*), whereas inclusion of *Wilaru* results in *Vegavis* being placed as the sister taxon of a clade containing *Telmabates*(*Presbyornis*(*Conflicto+Anatalavis*). When *Wilaru* is included, bootstrap support scores rise to 99% per node within the group *Presbyornis*(*Anatalavis+Conflicto*). Additional topology changes when *Wilaru* is included in the morphological analyses include *Dendrocygna guttata* being placed as the sister taxon of *Thalassornis leuconotus* and differences in the placements of several additional crown anatids (*Mergus serrator*, *Netta rufina*, *Cygnus atratus* and *Coscoroba coscoroba*; see Supplementary Data).

Within the combined results, addition of *Wilaru* further defines the base of included Galliformes recovered in the morphological analyses (*Gallinuloides wyomingensis*(*Macrocephalon maleo*+*Megapodius freycinet*)) and collapses the base of the stem anatid clade into a polytomy that includes *Wilaru* (94% clade credibility; see Supplementary Data). Both combined results recovered a *Presbyornis*(*Anatalavis+Conflicto*) clade, with posterior probability values for this clade again rising when *Wilaru* is included. When *Wilaru* is excluded, *Vegavis* is placed as the sister taxon to a *Telmabates*(*Presbyornis*(*Anatalavis+Conflicto*)) group as in the morphological results with *Wilaru* included, although the clade credibility of this recovered group is low (85% or less, see Figure 5 and Supplementary Data). These drastically different placements across the combined and morphological data analyses are likely due to the highly fragmentary nature of *Wilaru* specimens, although it does suggest that *Wilaru* may not be a presbyornithid as previously hypothesized (De Pietri et al. 2016). Due to this, we will largely focus on analyses that exclude *Wilaru*.

The resulting tree from the Bayesian analysis of the combined data matrix with *Wilaru* excluded are presented in Figure 5. Placement of extant taxa in the combined results are consistent with the resulting phylogram of Sun et al. (2017) based on mitochondrial genomes; however, this tree included relatively few anseriform taxa. Our Bayesian analysis results remain largely consistent with those of the phylogram of Sun et al. (2017) based on two mitochondrial genes, which contains almost all extant anseriform taxa used in our analysis. Differences arise in the placement of several non-goose taxa within crown Anatidae. We recover an a clade containing *Oxyura*+*Stictonetta* (*Anas*+*Amazonetta*(*Netta*(*Mergus*(*Tadorna*+*Chloephaga*)))) as being sister to the goose clade *Cygnus*+*Coscoroba*(*Branta*(*Anser+Anser*)), whereas Sun et al. 2017 recovers a *Mergus*(*Tadorna+Chloephaga*(*Netta*(*Anas*+*Amazonetta*))) group that is sister to *Oxyura*+the goose clade. Differences between these analyses are likely due to our inclusion of morphological data, mitochondrial genomes for more taxa and nuclear genes. Additionally, the Sun et al. (2017) analysis did not include *Stictonetta*, *Coscoroba*, or *Thalassiornis*, and our analysis did not include several genera included in Sun et al. (2017) (*eg. Neochen*, *Melanitta*, *Aythya* and more).

Within the combined results (Figure 5), clade credibility values of all nodes were 94% or higher with the exception of the placement of *Asteriornis maastrichtensis* as the sister taxon of a tinamou+lithornithid clade (54%), placement of *Crax* as the sister taxon to a *Lophura*+*Gallus gallus* sister group (69%), placement of *Paakniwatavis grandei* as the sister taxon to *Anseranas semipalmata*+(a fossil clade+crown Anatidae) (85%), placement of *Anseranas* as the sister taxon to a stem anatid clade and crown Anatidae (89%), placement of *Telmabates* as the sister taxon to *Prebyornis*(*Conflicto*+*Anatalavis*) (59%), placement of *Presbyornis* (55%), placement of *Anatalavis*+*Conflicto* (88%), and placement of *Netta rufina* as the sister taxon to a clade containing *Mergus*(*Tadorna+Chloephaga*) (85%).

The resulting strict consensus tree from analysis of morphological data is broadly confluent with combined results but presents alternative hypotheses for placement of several extinct and extant taxa within stem and crown Anatidae (see Supplementary Data).

Unconstrained parsimony analysis of morphological data resulted in two most parsimonious trees (MPTs) of 2,821 steps (CI = 0.291, RI = 0.580, RC = 0.169, HI = 0.709; see Supplementary Data). Morphological analysis recovers a fossil clade of stem Anatids containing *Telmabates+Presbyornis*(*Vegavis*(*Conflicto+Anatalavis*)) as the sister taxon to Anatidae, whereas this fossil clade containing the same taxa is structured as follows within the combined data results: *Vegavis*(*Telmabates*(*Presbyornis*(*Conflicto*+*Anatalavis*))). The positions of several extant anatids differ across the analyses as well. Within the morphological results, *Dendrocygna* is sister to *Thalassiornis*+crown Anatidae, whereas *Dendrocygna* and *Thalassiornis* are sister taxa in the combined data results. Additional differences within the morphological results comprise the placements of *Netta*, *Oxyura*, *Stictonetta*, the *Chloephaga*+*Tadorna* sister group, and *Mergus*. *Cygnus* and *Coscoroba* are additionally no longer sister taxa in the morphological results.

Both the combined and morphological analyses recover a *Diatryma*(*Pelagornis*+*Protodontopteryx*) stem anseriform clade (<50% bootstrap support, 99% clade credibility) and a clade as sister taxon to Anatidae containing *Telmabates*, *Presbyornis*, *Vegavis*, *Conflicto*, and *Anatalavis* (<50% bootstrap support, 95% clade credibility) despite the large number of morphological characters scored for these taxa and extensive reassessment of *Presbyornis* material. Recovery of a clade containing solely extinct taxa here may be real or discovered to be a paraphyletic assemblage; all synapomorphies for this clade exhibited CI**≤**0.5 in the morphological results and CI**≤**0.5 in the combined data results (see Supplementary Data). Our results for *Diatryma* and the Pelagornithidae are consistent with those of Bourdon (2005), in which Pelagornithidae are the sister taxon of Anseriformes; however, this study did not recover a monophyletic Galloanseres. Field et al. (2020) recovered the Pelagornithidae either as the sister taxon to an Anseriformes(*Conflicto*+*Anatalavis*) group or as the sister taxon of Charadriiformes.

Mayr (2011) recovered Pelagornithidae as the sister taxon to a Sylviornithidae(Dromornithidae(Galloanseres))) group, and Mayr et al. (2021) recovered a polytomy comprising Pelagornithidae, Galloanseres and Neoaves.

*Asteriornis* is recovered as a stem Galloanserine in the morphology only topology (<50% bootstrap support), but is the sister taxon to a tinamou+lithornithid clade within the combined results (54% clade credibility). Placement of *Presbyornis*, *Anatalavis*, and *Vegavis* as stem- Anatids is consistent with the results of Ericson (1997), Livezey (1997), and Elzanowski and Stidham (2010) for *Presbyornis* and both *Vegavis* and *Presbyornis* in the Livezey (1997) matrix (Clarke et al., 2005). Worthy et al. (2017) recovered *Presbyornis* and *Wilaru* as either the sister group of *Anseranas*+Anatidae or *Anseranas*, and recover *Vegavis* as either the sister taxon of anseriform-like Gastornithiformes (terror birds such as *Gastornis*) or the sister taxon to Anseriformes. Tambussi et al. (2019), utilizing the Worthy et al. (2017) data matrix, similarly recover *Presbyornis*, *Wilaru*, *Anatalavis* and *Conflicto* as stem Anseriformes and place *Vegavis* as the sister taxon of Gastornithiformes within stem Anseriformes. Field et al. (2020) recover *Wilaru*, *Presbyornis*, *Conflicto*, and *Anatalavis* as stem Anseriformes outside Anhimidae+Anatidae or *Wilaru* and *Presbyornis* as the sister group of *Anseranas.* Field et al (2020) also recovered *Vegavis* as a stem Galloanserine or Neoavian taxon (it remains in an unresolved polytomy) or as a stem Neornithine, and place *Asteriornis* as a stem Galloanserine or stem galliform. Torres et al. (2021) places *Conflicto* and *Vegavis* within a polytomy containing Anatidae and recovers *Asteriornis* as the sister taxon of a *Lithornis*+tinamou clade.

Within the combined data results, 7 unambiguous and 3 ambiguous optimized synapomorphies of the quadrate, coracoid, scapula, humerus, femur, tibiotarsus, tarsometatarsus, and pedal phalanges (two with CI = 1) support placement of *P. grandei* within Anseriformes.

Within the quadrate, the ventral apex of the crista tympanica terminates within the ventral half of the quadrate (character 172: state 2, ambiguous). The labrum externa along the lateral angle of the sternal coracoid is ventrocranially angled (416:2, unambiguous). Within the scapula, the acromion process is truncate and does not reach cranially beyond the articular faces of the head (437:1, ambiguous). The tuberculum ventral and crista along the proximal margin of the fossa pneumotricipitalis are domed and distally prominent, overhanging the fossa pneumotricipitalis (456:2, unambiguous, CI = 1). The length of the femur is approximately one half the length of the tibiotarsus (597:1, unambiguous). The distal opening of the pons supratendinous of the tibiotarsus is centered along the midline (653:3, unambiguous). Within the tarsometatarsus the medial margin of the medial cotyle is exceptionally projected proximally and crista-like (658:2, unambiguous, CI = 1), the lateral cotyle is flattened or only slightly concave (662:2, ambiguous), and the hypotarsal eminence is proximally prominent (664:2, unambiguous). Within the pedal phalanges, pedal phalanx II: digit 2 is slightly more elongate than III:2 (706:2, unambiguous).

Seven unambiguous and three ambiguous optimized synapomorphies of the quadrate, mandible, furcula, scapula, carpometacarpus, tarsometatarsus and pedal phalanges (one with CI = 1) support placement of *P. grandei* as a stem anseranatid. Within the quadrate, the crista tympanica is a present and extremely prominent crista (171:3, unambiguous) and the caudal face of the otic process is deeply concave (176:2, unambiguous). Within the mandible, the medial portion of the ramus caudal to the coronoid process (or homologous site) is extremely deep and concave (244:2, unambiguous), and a true retroarticular process is present and exceptionally tapered throughout (256:2, ambiguous). The width of the lateral diameter of the furcular ramus is larger than that at the symphysis (436:3, ambiguous), and the scapula is shorter than the humerus in length (449:1, unambiguous). Within the carpometacarpus, the proximal terminus of the dorsal rim of the trochlea carpalis is strongly angular and proximally elongated (514:2, unambiguous). Within the tarsometatarsus, the distal-most terminus(i) of medial crest(s) are much more distally extensive and the lateral crista(e) are proximodistally truncate, about 1/2-2/3 proximodistal length of the medial crista (669:1, unambiguous, CI = 1). The proximal portion of the sulcus extensorius medial to the dorsal foramina vascularia proximalia is present and deeply excavated (680:2, ambiguous). Within the pedal phalanges, phalanx III is longer than the tarsometatarsus (719:1, unambiguous).

Ancestral state reconstruction for behavioral characters using the combined results phylogeny suggests that *P. grandei* likely preferred an aquatic or semi-aquatic environment, was either not specialized for aquatic feeding modes (hereafter referred to as a “non-swimmer”) or was a surface swimmer, was primarily herbivorous, was a grazer, had some form of rhamphothecal lamellae, and that the length of its pedal digit III was subequal to or longer than that of the tarsometatarsus (confirmed through scoring), had an ossified pessulus of the syrinx, and did not have asymmetry at the tracheobronchial juncture of the syrinx. Status of pedal webbing could not be reconstructed for *P. grandei*; however, the anatomy and skeletal proportions of *P. grandei* indicate an aquatic surface swimmer. This is especially likely as *P. grandei* has a pedal digit III that is longer than the tarsometatarsus, indicating that this taxon likely was a surface swimmer and led an aquatic or semi-aquatic lifestyle (Manegold 2006; Birn- Jeffrey et al. 2012). Anseriformes including stem-Anseriformes (*Diatryma*+Pelagornithidae) preferred a terrestrial or aquatic habitat, were non-swimmers, were omnivorous or carnivorous, were mixed feeders or grazers, did not have pedal webbing, had a pedal digit III was shorter than the length of the tarsometatarsus, had an ossified pessulus, and had no asymmetry at the tracheobronchial juncture. The status of rhamphothecal lamellae could not be reconstructed for this clade. Crown Anseriformes ancestrally were identical with the exception of preferring an aquatic or semiaquatic habitat, being primarily herbivorous, having a form of rhamphothecal lamellae, and being grazers. Anatidae including the fossil clade of stem Anatids preferred an aquatic or semiaquatic habitat, were surface swimmers, were primarily herbivorous, had full rhamphothecal lamellae present, were grazers, possessed pedal webbing, had a pedal digit III was subequal in length or longer than the tarsometatarsus, had an ossified pessulus, and had no asymmetry at the tracheobronchial juncture. Crown Anatids were ancestrally identical with the exception of preferring only aquatic environments.

Ancestral state reconstruction for behavioral characters using the morphological results provided identical results for *P. grandei.* Status of pedal webbing again could not be reconstructed for *P. grandei.* Ancestral state reconstructions for Anseriformes including stem- Anseriformes (*Diatryma*+Pelagornithidae) were identical to those of the combined data analysis. Again, the status of rhamphothecal lamellae could not be reconstructed for this clade. Ancestral state reconstructions for crown Anseriformes remained identical. Reconstructions again remained identical for Anatidae including stem Anatids, with the exception of asymmetry at the tracheobronchial junction being unable to be reconstructed. Reconstructions for crown Anatidae were identical to those recovered using the combined data results, again with the exception of asymmetry at the tracheobronchial junction being unable to be reconstructed.

PCA and LDA analyses included all 32 skeletal measurements of the Hinic-Frlog and Motani (2010) dataset which included aquatic taxa and behaviors of diverse avian subclades; however, 12 measurements maximum could be completed per extinct taxon, with most including less. Results were largely not illuminating with respect to the new fossil. When the PCA analysis was re-run using only seven skeletal measurements that had been identified by Hinic-Frlog and Motani (2010) as best describing swimming mode, resulting proposed swimming modes for these taxa were identical (see Supplementary Data). *Presbyornis* was recovered as a non-swimmer and *Vegavis* and *Anatalavis* as wing-propelled divers or surface swimmers with the extant anhimid *Chauna* as either a wing-propelled diver or non-swimmer. *P. grandei* and *Telmabates* were recovered as foot-propelled divers or surface swimmers (see Supplementary Data). Linear discriminant analysis of the same dataset predicted that *P. grandei*, *Telmabates* and *Chauna* were surface swimmers, *Anatalavis* as a plunge diver, as wella s *Presbyornis* and *Vegavis* as foot-propelled divers (see Supplementary Data). While the PCA results but not the LDA results for *Presbyornis* are consistent with its skeletal proportions, the rest of the results are largely inconsistent with other features or the known behaviors (i.e., *Chauna*) of the rest of the assessed taxa. New analyses tailored to the behaviors seen within Anseriformes would likely be needed to usefully adapt the comprehensive dataset of Hinic-Frlog and Motani (2010).

## Discussion

We recover *Paakniwatavis grandei* as the sister taxon to an *Anseranas*+Anatidae clade within Anseriformes across all analyses, regardless of taxon sampling. This placement is consistent with the unique combination of anhimid-like and anseranatid-like morphologies displayed by the taxon as well as its aquatic morphologies. A *Diatryma*+Pelagornithidae group was also recovered across all analyses as the sister taxon of crown Anseriformes, and a clade containing *Vegavis iaai*, *Prebyornis*, *Telmabates*, *Conflicto*, and *Anatalavis* was recovered as the sister taxon of crown Anatidae across all analyses. Recovery of additional, more complete and better preserved anseriform-like fossils is necessary to more robustly resolve the phylogenetic placement of these important taxa; however, consistent results across several analyses using different data types and methods suggests that these placements are fairly robust. *P. grandei* represents a unique ecology for the Green River Formation, and this new fossil along with this new dataset and re-evaluation of *Presbyornis* material thus begins to elucidate several critical issues in anseriform evolution.

Ancestral state reconstruction across all analyses suggests the evolution of a combination of terrestrial and semi-aquatic traits at the base of Anseriformes, with crown Anseriformes exhibiting a shift toward more aquatic traits such as preferring an aquatic or semiaquatic habitat, being primarily herbivorous, having a form of rhamphothecal lamellae, and being grazers (Figure 5). This is inconsistent with the assertion that Anseriformes were ancestrally terrestrial as suggested by Olson and Feduccia (1980), Ericson (1997), and Livezey (1997). The placement of *P. grandei* and its influence on the optimization of these reconstructions suggests aquatic or semi-aquatic ancestry of Anseriformes, especially combined with a reduced form of rhamphothecal lamellae present in extant Anhimidae (Olson and Feduccia, 1980).

The question of how anseriform beak morphology and filter feeding evolved remains an open one. Either a narrower, “goose-like” beak was ancestral for Anseriformes, or this narrower beak evolved several times within Anseriformes (Olsen, 2017). The “goose-like” beak is associated with increased leaf consumption, decreased invertebrate consumption, and an increase in the mechanical advantage of the beak that allows for more effective cropping of plants (Olsen, 2017), whereas “duck-like” beaks are associated with increased filter feeding and consumption of invertebrates. We take this classification of a “goose-like” beak a step further in the broader context of both stem and crown Anseriformes: We first identify whether the rostrum is mediolaterally wider than the width of the paroccipital processes (indicating an anseranatid-like beak; character 3) and, if so, whether it is anteriorly tapered and narrowed further (a “goose-like” beak; character 4: state 1), or whether it remains subequal in width (a “duck-like” beak; 4:2). *P. grandei* is the only Paleogene anseriform known to present a narrow, anhimid-like beak other than the Pelagornithidae. Its beak is mediolaterally narrower than the width of its paroccipital processes, as in extant Anhimidae. *Chaunoides* and *Telmabates* have no preserved skull and *Vegavis* has no preserved rostrum or braincase, while other Paleogene fossils with a beak preserved such as *Presbyornis*, *Anatalavis* and *Conflicto* all present an anseranatid-like beak. All of our analyses posit the first appearance of a beak that is mediolaterally wider than the width of the paroccipital processes as ancestral to the node containing *Anseranas semipalmata*. It would have been an anteriorly tapered, “goose-like” beak like that of the extant Anhmidae or *Anseranas* (Figure 5). Across all analyses, ancestral state reconstruction indicates that this “goose-like” beak is ancestral to Anseranatidae+Anatidae. All analyses indicate that anteriorly wider “duck-like” beaks evolve at least twice: once after the divergence of *Anseranas* in taxa closer to Anatidae (present in *Presbyornis* and *Anatalavis*), and once within crown Anatidae (see Figure 5 and Supplementary Data). These results contradict those of Olsen (2017), who found that a duck-like beak was ancestral for most Anatidae, followed by multiple transitions toward a goose-like beak; however, this study performed ancestral state reconstruction using a phylomorphospace of beak curvature measurements, beak function metrics, and quantified diet data for a smaller sample of anseriform taxa that included only two extinct taxa, *Presbyornis* and the recently extinct moa- nalo *Thambetochen chauliodous* Olson and Wetmore 1976 (Olson and James, 1991).

At the same time, our results are consistent with the results of Olsen (2017) in that we find rhamphothecal lamellae to have been present ancestrally for crown Anseriformes and Anatidae (including within both the stem and crown lineages of these groups), indicating that herbivory and/or filter feeding was ancestral for these clades. This again contradicts the hypothesis that Anseriformes were ancestrally terrestrial and would explain the presence of reduced rhamphothecal lamellae in extant Anhimidae (Olson and Feduccia, 1980). Anhimidae represent the only known example of rhamphothecal lamellae being present without pedal webbing in extant birds; however, similar lamellae-like ridges have been found in Ornithomimus (Norell et al., 2001; Barrett, 2005), Gallimimus, chelonians, hadrosaurs (Barrett, 2005) and an edentulous ceratosaur (Alves de Souza et al., 2021). A partial correlation between the presence of these ridges and exclusively herbivorous diet among terrestrial chelonians has been found (Bramble, 1974; Pritchard, 1979), suggesting that more prominent ridges were present when more coarse vegetation was eaten. Studies on the jaw mechanics, locomotion and gut contents of hadrosaurs have similarly demonstrated that they were obligate terrestrial herbivores that used their beak for cropping through vegetation (Weishampel, 1984; Forster, 1997). If our ancestral state reconstructions for crown Anseriformes are correct, lamellae coevolved with a shift toward herbivory and grazing along with the preference for a more aquatic habitat despite a lack of pedal webbing. Based on the available evidence and our results, some form of rhamphothecal lamellae was ancestral to crown Anseriformes and could have developed due to aquatic grazing, then remained (or became reduced) within extant Anhimidae while developing further within more derived crown anseriform taxa. Webbing then was maximally ancestral to *P. grandei*, and minimally was ancestral to *Anseranas*. Our results thus suggest that crown Anseriformes were maximally ancestrally aquatic or semi-aquatic, and fully aquatic ancestral to crown Anseranatidae. In general our results suggest a trend within Anseriformes toward aquatic grazing and the anseranatid “goose-like” beak to obtain an herbivorous (increased leaf and root consumption) diet, with at least two evolutions of a “duck-like” beak associated with increased filter feeding and invertebrates obtained by this feeding mode at least once within stem-Anatidae and once within Anatidae (Kooloos et al., 1989; Van der Leeuw et al., 2003). If these placements and ancestral state reconstructions are correct, this would add to mounting support that feeding ecology has acted as the primary selective force in waterfowl beak shape diversification (Olsen, 2017).

Further elucidation of anseriform behavioral evolution is indicated in ancestral state reconstruction of syringeal characters. Ancestral state reconstruction within the combined results indicates that *P. grandei*, Anseriformes (inclusive and exclusive of stem Anseriformes) and Anatids (inclusive and exclusive of stem Anatids) had an ossified pessulus of the syrinx, a derived neognath bird feature that has been proposed to anchor enlarged vocal folds or labia (King, 1989), consistent with the results of Clarke et al., (2016). These taxa also were indicated to ancestrally not possess asymmetry at the tracheobronchial juncture of the syrinx. Ancestral reconstruction within the morphological results is identical with the exception of ambiguity within stem and crown Anatidae regarding asymmetry at the tracheobronchial juncture. Both results indicate that asymmetry at the tracheobronchial juncture must have evolved at least once within Anseranatidae+Anatidae. This is somewhat consistent with Clarke et al. (2016); however, while Clarke et al. (2016) considered a single origin in Anatidae likely, our results may suggest more gains and losses within this Anseranatidae+Anatidae clade. Further study and coding of extant syrinx asymmetry is necessary as previous descriptions and figures of extant syrinxes largely focus on pronounced asymmetrical bullae in some male anseriform taxa rather than the more subtle asymmetry of the rings found in females, and the large range of variation across differing taxa and sexes within Anseriformes is not well understood (Johnsgard 1962; King 1989). Better understanding asymmetry in extant taxa has important implications for extinct taxa as well; for example, the subtle asymmetry present in *Vegavis* may suggest that this taxon exhibited sexual dimorphism within the syrinx as in some extant Anatidae. Asymmetry is an important trait to further study as it is correlated with the presence of a dual sound source and the presence of labia (King, 1989; Clarke et al., 2016). Recovery of further fossils that include syrinxes and further study of extant syrinx anatomy and function in extant birds is thus necessary to understand the evolution of this organ.

All analyses recover the Cretaceous-Paleogene taxa *Vegavis*, *Anatalavis*, *Conflicto*, *Presbyornis* and *Telmabates* within a clade that is the sister taxon to Anatidae. Within the combined data results, six unambiguous and 10 ambiguous synapomorphies (CI < 1) were recovered for this clade that united three or more of these taxa within the axial skeleton and hindlimbs (see Supplementary Data). Although several appear to be plesiomorphic, these characters may represent evolutionary and biological/ecological significance pending recovery of key elements from taxa with more missing data such as *Vegavis*, *Telmabates* and *Conflicto*.

Although this character was not optimized as a synapomorphy for this clade, *Anatalavis*, *Temabates*, *Presbyornis*, and the Neogene Dromornithidae have amphicoelous thoracic vertebrae (Martin 1987; Ericson, 1997; Clarke, 2004), whereas *Vegavis*, Pelagornithidae and gastornithids have heterocoelous thoracic vertebrae (Clarke et al. 2005, 2016). Although it cannot be discerned which non-heterocoelous form the vertebrae of *P. grandei* possess due to taphonomic distortion, it is likely that *P. grandei* possessed amphicoelous vertebrae as well. Amphicoelous thoracic vertebrae are plesiomorphic within Avialae but also present in well-nested neoavian clade Charadriiformes (Martin, 1987; Ericson, 1997; Mayr and Clarke, 2003; Clarke, 2004).

Opisthocoelous thoracic vertebrae are only known within Neoaves (Mayr and Clarke, 2003; Livezey and Zusi, 2006; Musser and Cracraft, 2019), and amphiplatyan and procoelous vertebrae are only known in non-avian dinosaurs (Livezey and Zusi, 2006).

Further study on the function of amphicoelous vertebrae in the context of avian evolution and ecology is needed, especially since birds possess a unique dorsal intervertebral joint (Wintrich et al., 2020); however, the literature on this in fishes and crocodylomorphs suggests that amphicoelous vertebrae provide a more rigid spine (Yakovlev, 1967; Laerm, 1976; Molnar et al., 2015) that can withstand increased stress without deformation that may be caused by powerful movements of musculature (Yakovlev, 1967; Laerm, 1976), allowing for rapid flexure of the spine. Amphicoelous vertebrae have evolved several times in sharks, dipnoans, bony ganoids and teleosts, associating their appearance with improved speed of motion (Yakovlev, 1967).

Amphicoelous vertebrae are also associated with aquatic environments, as transitions from amphicoelous to platycoelous vertebrae has been hypothesized to represent a transition from aquatic to terrestrial environments (Romer, 1956).

Results further clarify the complex picture of avian evolution around the K-T boundary, indicating that several lineages within Anseriformes with a variety of ecologies not represented in the crown were present by the latest Cretaceous and into the early Paleogene. *P. grandei* represents an early Eocene lacustrine, aquatic taxon that swam and likely used its narrow bill in aquatic grazing or mixed feeding. The Cretaceous-early Paleogene *Anatalavis* and *Presbyornis* were aquatic taxa that likely filter fed on a more invertebrate-heavy diet within both marine and marine and lacustrine environments, respectively (Olsen, 2017). The early Eocene *Telmabates* may have shared a similar ecology to *Presbyornis* given its amphicoelous thoracic vertebrae and evidence that the Casamayor formation in which it was found is known to be a marine-fluvial transition zone (Raigemborn et al., 2010). Other Cretacous and Paleogene material has been referred to *Presbyornis* from more fluvial as well as marine settings possibly suggesting a cosmopolitan, flexible habitus (Olson, 1994; Clarke and Norell, 2004; Kurochkin and Dyke, 2010; Hood et al., 2019). At the same time the Cretaceous *Vegavis*, with heterocoelous thoracic vertebrae, was present in a near shore marine environment with unknown diet, and the Cretaceous *Conflicto* was present within a near shore marine/transitory estuarine environment, again with unknown diet and locomotion due to missing data (Tambussi et al., 2019). In addition to this array of taxa and ecologies, the specialized Cretaceous-Paleogene marine, piscivorous pseudodonts and the giant terrestrial Paleogene Gastornithids were present within the stem anseriform lineage. If *Vegavis* and *Conflicto* are also found to have had omnivorous, mixed and/or piscivorous diets, a proliferation of Creataceous-early Paleogene non-herbivorous stem and crown Anseriformes may have arisen. Proposed significant loss of plant cover due to global cooling and the K-T impact event (Crane and Lidgard, 1989; Kaiho et al., 2016; Field et al., 2018; Condamine et al., 2020; Lyons et al., 2020; Li et al., 2021) could suggest a short but strong selective regime favoring mixed and non-herbivore specialists. Fossil evidence suggests that early Anseriformes were diversifying rapidly since at least the late Cretaceous and were already widespread within the same time frame, as the early Eocene *Telmabates* was found in Patagonia and Paleocene and Eocene *Presbyornis* material has been recovered from North America, Europe and Mongolia (Olson, 1994; Clarke and Norell, 2004; Kurochkin and Dyke, 2010; Grande, 2013; Hood et al., 2019).

An approximate mean body mass estimate for *P. grandei* is 304.4g based on the published allometric equation using femoral length from Field et al., 2013. This body mass level is quite small; it is most comparable to the mass of many *Anas* (within the 300s range). Many other anatids are generally larger (which typically range from 600-over 1,000g; Dunning, 2007). Its body mass is estimated to be less than half that of *Presbyornis* and *Telmabates* (882.2g and 1423.0g, respectively). This is consistent with recent evidence that correlation between herbivory and body mass is not significant when accounting for phylogeny (Olsen, 2015).

Further analysis of these Paleogene anseriform fossils in the context of broader extinct taxon sampling, especially in the context of extinct anseriform-like taxa, is necessary to further evidence placement of *P. grandei* and other Paleogene anseriform-like taxa and to gain more robust insight into ancestral states; however, *P. grandei* represents a key taxon, a unique ecology within known Anseriformes and the Green River Formation, and a potential calibration point for anseranatids. Other included extinct taxa also represent potential calibration points that would be valuable for stem Anseriformes, Anhimidae and stem Anatidae, although recovery of additional fossils and further phylogenetic analyses (especially of taxa such as *Chaunoides* and *Wilaru*) are preferable to confirm these relationships, further reveal the ecological and behavioral evolution and biogeography of Anseriformes, and better elucidate our understanding of avian evolution.

## Author Contributions

Conceptualization, all authors; Methodology, G.M. and J.A.C.; Software, G.M.; Validation, all authors; Formal Analyses, G.M. and J.A.C.; Investigation, all authors; Resources, all authors; Data Curation, G.M.; Writing—Original Draft Preparation, G.M.; Writing—Review and Editing, all authors; Visualization, G.M. and J.A.C.; Supervision, G.M. and J.A.C.; Project Administration, G.M. and J.A.C.; Funding Acquisition, G.M.

## Acknowledgments

We thank all of the staff of FMNH, especially Lance Grande, Jingmai O’Connor, William Simpson, Adrienne Stroup, Shannon Hackett, John Bates, and Ben Marks for specimen access and valuable discussion. We thank April Neander for scanning the specimen, and Matthew Colbert for aid in scan visualization and segmentation. We thank Helen James, Christopher Milensky, Brian Schmidt and Mark Florence of the Smithsonian National Museum of Natural History for access to the Ornithology and Vertebrate Paleontology collections. We thank Joel Cracraft, Paul Sweet, Mark Norell, Ruth O’Leary and Carl Mehling of AMNH for access to the Ornithology and Vertebrate Paleontology Collections and thank the Ornithology Department of AMNH for the Ornithology Collections Study Grant that aided in funding this work. We additionally thank Kenneth Bader, Matthew Brown, and Christopher Sagebiel for access to additional TMM collections and their aid in working with AMNH loans. Finally, we thank Joel Cracraft, Zhiheng Li, Melissa Kemp, Christopher Bell, Daniel Field, Daniel Ksepka and Christopher Torres for valuable discussion.

## Data Availability Statement and Supplementary Material

The datasets generated and analyzed for this study as well as supplementary material can be found in Morphobank (O’Leary and Kaufman 2012) under Project 4001 (http://morphobank.org/permalink/?P4001) and character descriptions can be found in the Appendix.

